# Higher eukaryote-specific APC/C composition is a determinant of spindle assembly checkpoint importance

**DOI:** 10.1101/222422

**Authors:** Thomas Wild, Magda Budzowska, Gopal Karemore, Chunaram Choudhary

## Abstract

The multisubunit ubiquitin ligase APC/C (anaphase promoting complex/cyclosome) is essential for mitosis by promoting timely degradation of cyclin B1. Proper timing of APC/C activation is regulated by the spindle assembly checkpoint (SAC), which is initiated by the kinase MPS1 and implemented by MAD2-dependent inhibition of the APC/C. Here we analysed the contribution of the higher eukaryote-specific APC/C subunits APC7 and APC16 to APC/C composition, function and regulation. APC16 is required for APC7 assembly into the APC/C, while APC16 assembles independently of APC7. ΔAPC7 and ΔAPC16 cells display no major defects in mitotic progression, cyclin B1 degradation or SAC response. Strikingly, however, deletion of either APC7 or APC16 is sufficient to provide synthetic viability to MAD2 deletion. ΔAPC7ΔMAD2 cells display an accelerated mitosis and require SAC-independent MPS1 function for maintaining their genome stability. Overall, these results show how human APC/C composition critically influences the cellular fate upon loss of SAC activity.

## Introduction

Eukaryotic cells depend on the activity of the anaphase promoting complex / cyclosome (APC/C), a multi-subunit ubiquitin ligase, to progress through mitosis (Peters 2006, Pines 2011, Primorac and Musacchio 2013). Among a variety of mitotic APC/C substrates, securin and CCNB1 are the only essential APC/C substrates for mitotic progression in yeast (Thornton and Toczyski 2003). The APC/C requires the co-activator CDC20 as well as an E2 enzyme, UBE2C or UBE2D, to initiate CCNB1 ubiquitylation (Hershko et al. 1994, Yu et al. 1996, Townsley et al. 1997, Wild et al. 2016). Substrate-conjugated ubiquitin chains can be elongated by the APC/C with the E2 enzyme UBE2S, enhancing substrate degradation efficiency (Garnett et al. 2009, Williamson et al. 2009, Wu et al. 2010, Meyer and Rape 2014, Min et al. 2015). Lowering APC/C activity delays mitotic progression, whereas loss of APC/C activity is lethal.

The timing of APC/C-initiated securin and CCNB1 degradation is of critical importance for the faithful segregation of chromosomes. Proper timing is achieved by the spindle assembly checkpoint (SAC), which coordinates bipolar chromosome attachment to the mitotic spindle with APC/C-mediated securin and CCNB1 ubiquitylation (London and Biggins 2014, Sivakumar and Gorbsky 2015). The SAC is activated by the recruitment of the kinase MPS1 to unattached kinetochores. Subsequent MPS1-catalyzed phosphorylation events recruit SAC proteins to unattached kinetochores, resulting in the formation of the mitotic checkpoint complex (MCC), consisting of MAD2, CDC20, BUBR1 and BUB3 (Sudakin et al. 2001). MCC binding to the APC/C blocks substrate recognition by CDC20 and hampers access of UBE2C to initiate substrate ubiquitylation (Alfieri et al. 2016, Yamaguchi et al. 2016). While SAC function is conserved from yeast to human, its activity apparently gained importance in higher eukaryotes: deletion of SAC proteins such as MAD2 is compatible with viability in yeast, yet leads to genomic instability in mice and human cells incompatible with life (Dobles et al. 2000, Michel et al 2001, Michel et al. 2004, Meraldi et al. 2004, Kops et al. 2005).

The APC/C consists of 19 subunits composed of 14 distinct proteins. Initial low resolution structures divided the global APC/C architecture into ‘platform’ and ‘arc lamp’ (also known as ‘TPR lobe’) (Dube et al. 2005). Recently, the atomic architecture of APC/C has been elucidated by cryo-electron microscopy (Chang et al. 2015). The two largest APC/C subunits, APC1 and APC2, form the core of the platform, while the TPR subunits APC3, APC6, APC7 and APC8 constitute the majority of the ‘arc lamp’. The catalytic center of the APC/C is formed by APC11 and APC2 along with APC10 and the co-activator CDC20 or CDH1 for substrate recognition. APC/C composition is conserved from yeast to human, except for two higher-eukaryote specific subunits, APC7 and APC16, located at the tip of the ‘arc lamp’ (Kops et al. 2010, Hubner et al. 2010, Hutchins et al. 2010, Ohta et al. 2011, Green et al. 2011, Herzog et al. 2009, Chang et al. 2015). APC7 is present in two copies and together with one APC16 molecule sits on top of APC3. Depletion experiments revealed a role of APC16 in mitotic progression and APC/C substrate stability, but not APC/C assembly (Kops et al. 2010, Shakes et al. 2011). Depletion of APC7 by RNAi in *Drosophila melanogaster* had limited impact on mitotic progression and, consistently, an APC7-null strain is viable (Pal et al. 2007). In vitro, it has been shown that APC3 forms a complex with APC7, and that this complex is stabilized by APC16 (Yamaguchi et al. 2015). Based on this data, the authors suggested that the APC3/APC7/APC16 subcomplex may constitute an assembly unit or that APC16 recruits APC7 onto APC3. Here we set out to analyse the contribution of APC7 and APC16 to APC/C assembly, function and regulation using genetic knockouts in human cells.

## Results

### APC16 is required for APC7 assembly into the APC/C in vivo

To study the impact of APC7 and APC16 on the composition of APC/C in vivo, we fused mCherry to the C-terminus of APC8 and investigated APC/C composition by affinity-purification mass spectrometry (AP-MS) (Poser et al. 2008). Because APC8 is essential for cell viability, biallelic tagging of the gene indicated functional integrity of APC8-mCherry-tagged APC/C. Next, we deleted APC7 or APC16 in the APC8-mCherry cell line (Figure S1A, Figure S1B) and analysed APC/C composition with SILAC-based quantitative AP-MS (Figure 1A). We purified the APC/C from wild-type and AAPC7 cells by APC8-mCherry affinity purification and compared the relative enrichment of APC/C subunits in the APC8-mCherry pulldowns over a mock pulldown. In wild-type cells, all APC/C subunits detected in the mass spectrometry analysis were enriched (>2-fold) from APC8-mCherry tagged cells compared to control (Figure 1B), demonstrating proper incorporation of APC8-mCherry into APC/C complexes. All APC/C subunits, except for APC7, co-purify with APC8-mCherry from AAPC7 cells, showing that the remaining APC/C subunits assemble in absence of APC7 (Figure 1B). We confirmed APC/C assembly in cells lacking APC7 by immunoblotting using antibodies for several APC/C subunits (Figure 1C). To analyse the role of APC16 in APC/C assembly, we used SILAC-based AP-MS to compare the APC/C composition in APC8-mCherry expressing wild-type and AAPC16 cells. Interestingly, APC/C enriched from AAPC16 cells not only lacked APC16, but also APC7 (Figure 1D, Figure S1C). To test whether expression of APC16 in AAPC16 cells restores APC7 incorporation into the APC/C, we expressed APC16-EGFP in AAPC16 cells and analysed APC/C composition (Figure 1E). Consistent with its integration into the APC/C (Kops et al. 2010), APC16-EGFP co-purified with APC8-mCherry and, indeed, restored APC7 co-purification with APC8-mCherry. Together, these results show that in vivo APC16 incorporates into the APC/C in the absence of APC7, whereas APC7 requires APC16 for incorporation into the APC/C.

**Figure 1.**
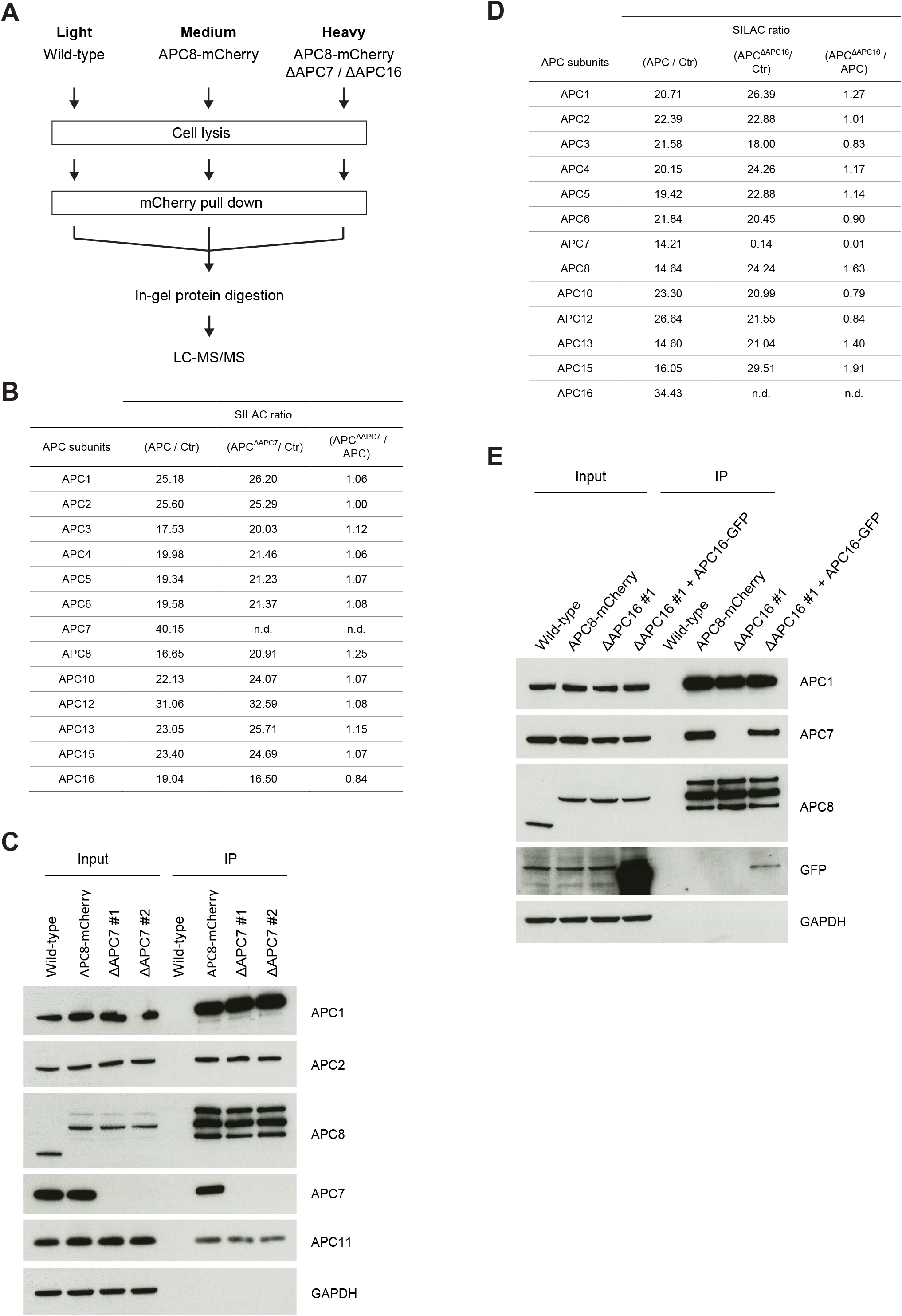
APC16 is required for the assembly of APC7 into the APC/C. A) Scheme of triple SILAC setup for MS-based analysis of APC/C composition. APC8-mCherry expressing wild-type, ΔAPC7, or ΔAPC16 cells were SILAC labelled as indicated and APC/C was purified using mCherry affinity beads. As a control, a mock pull-down was performed from light SILAC labelled cells. B) SILAC ratios for APC/C subunits enriched from APC8-mCherry wild-type and APC8-mCherry ΔAPC7 cells. APC/C was purified with mCherry pulldowns from wild-type (medium SILAC condition) and ΔAPC7 cells (heavy SILAC condition) and analysed by mass spectrometry. Mock pull-down from wild-type HCT116 cells served as control (light SILAC condition). The table shows combined SILAC ratios for all APC/C subunits detected in three technical replicates (n.d. = not determined). C) Analysis of APC/C composition in ΔAPC7 cells. APC/C was purified via APC8-mCherry pulldowns from APC8-mCherry and APC8-mCherry ΔAPC7 cells (from two independent ΔAPC7 clonal cell lines) and subsequently analyzed by immunoblotting using indicated antibodies. Wild-type cells were used as a control for unspecific binding to the affinity beads. GAPDH levels were analysed to verify equal amount of input for the different cell lines. D) SILAC ratios for APC/C subunits enriched from APC8-mCherry and APC8-mCherry ΔAPC16 cells. The analysis was performed as described in B). E) Immunoblot analysis of APC/C composition in ΔAPC16 cells with indicated antibodies. APC/C was purified via APC8-mCherry pulldowns from APC8-mCherry cells and from APC8-mCherry ΔAPC16 cells. Indicated samples were transiently transfected with APC16-EGFP 32 hours prior to the APC/C pulldown. Wild-type cells were used as a control for unspecific binding to the affinity beads.

### Cells lacking either APC7 or APC16 display no major defects in mitotic APC/C function

To analyse mitotic progression in cells lacking APC7 or APC16, we endogenously tagged histone H2B with mVenus and CCNB1 with mCerulean3 in wild-type, ΔAPC7 and ΔAPC16 background (Figure S2A, Figure S2B and Figure S2C) and used these cell lines to monitor mitotic timing by time-lapse microscopy. Analysis of mitotic timing (defined as the timing from nuclear CCNB1 influx to anaphase onset) revealed no significant difference between wild-type, ΔAPC7 and ΔAPC16 cells (Figure 2A, Figure S2C). Concordantly, analysis of mitotic CCNB1 degradation revealed no significant alteration in CCNB1 degradation kinetics in APC7 or APC16 knockout cells (Figure 2B). Next, we assessed the responsiveness of ΔAPC7 and ΔAPC16 cells to SAC activation. Cells were treated with nocodazole, which activates the SAC, for 18 hours and the cellular DNA content was subsequently analysed by flow cytometry (FACS) (Figure S2D). Nocodazole treatment caused a marked increase of 4N DNA containing cells for all analysed cell lines, showing that ΔAPC7 and ΔAPC16 cells remain sensitive to SAC signaling. To further verify SAC functionality in ΔAPC7 cells, we analysed the nocodazole-dependent association of SAC-generated MCC with the APC/C. For both wild-type and ΔAPC7 cells an increased enrichment of the SAC proteins - BUB3, BUBR1, CDC20 and MAD2 - was observed in APC8-mCherry pulldowns upon a nocodazole-induced mitotic arrest (Figure 2C and Figure 2D). To analyse the dependence of ΔAPC7 and ΔAPC16 cells on SAC function for genomic stability, we treated cell lines with the MPS1 inhibitor reversine and subsequently analysed cellular ploidy by FACS (Figure S2E). Reversine treatment resulted in an increased number of polyploid cells (>4N DNA content) for wild-type, ΔAPC7 and AAP16 cells, demonstrating the importance of MPS1 activity for maintaining genome stability in these cell lines. To independently assess the importance of SAC function in ΔAPC7 and ΔAPC16 cells, we depleted cells of MAD2 by RNAi and analysed nuclear morphology four days thereafter (Figure 2E and Figure S2F). Consistent with the crucial role of MAD2 in maintaining genomic stability, we quantified a marked decrease in normally sized and shaped nuclei in wild type cells upon MAD2-depletion (Figure 2F). Interestingly, however, the decrease in normal nuclei upon MAD2 RNAi was reduced in both ΔAPC7 and ΔAPC16 cells (Figure 2F).

**Figure 2.**
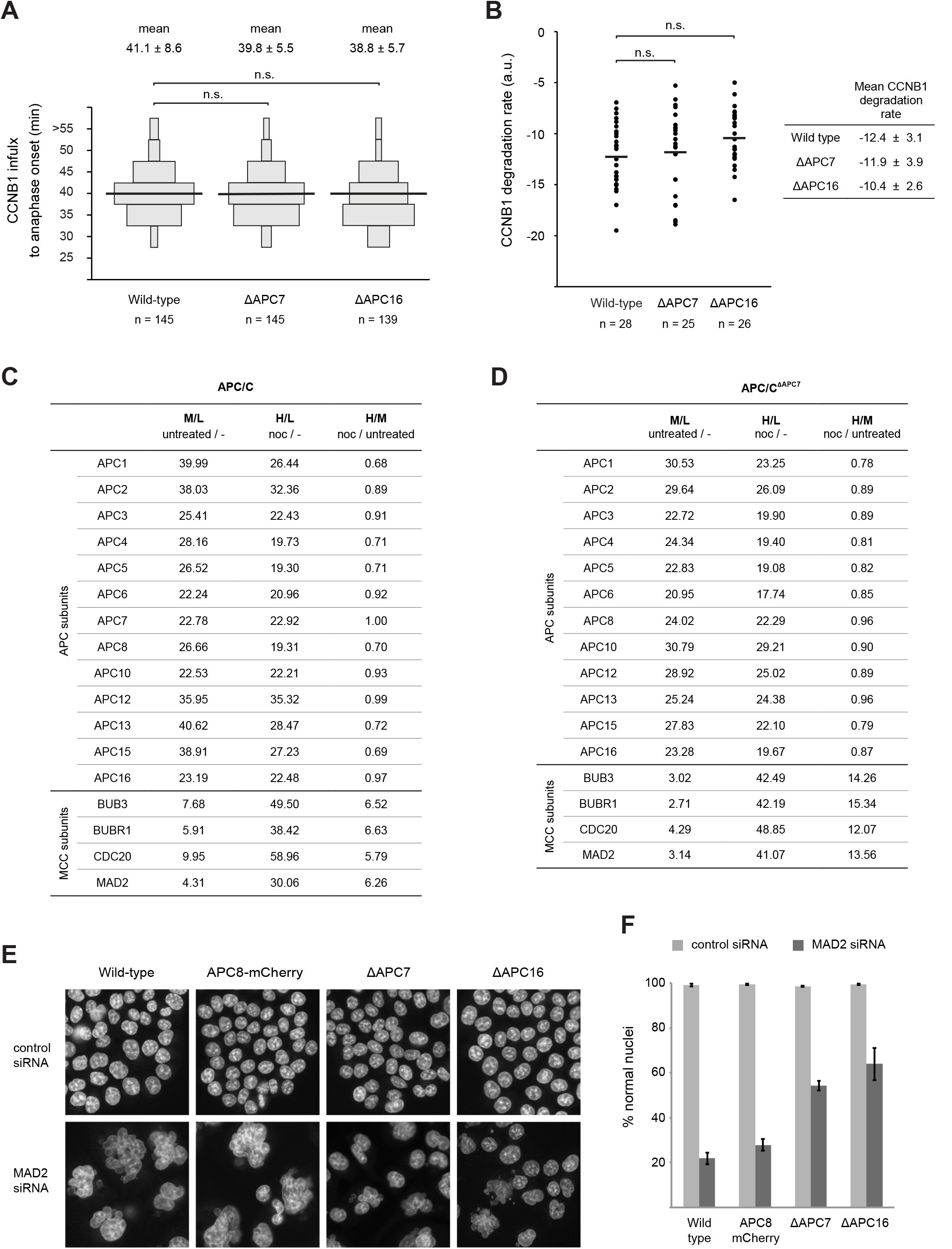
Analysis of mitotic APC/C function in ΔAPC7 and ΔAPC16 cells. A) Time between nuclear CCNB1 influx and anaphase onset for indicated cell lines. Wild-type, ΔAPC7 and ΔAPC16 cell lines expressing mVenus-tagged histone H2B and mCerulean3-tagged CCNB1 were imaged every 5 min. The data shows time from nuclear CCNB1 influx to anaphase onset for individual cells. Cells with identical timing are depicted as a box and the size of each box is scaled according to the percentage they contribute to the whole population of the respective cell line. The mean timing and the standard deviation are stated on top. The median timing is drawn as a black line. Results of three independent experiments are shown and the number (n) of analysed cells is stated for each cell line. A two-tailed t-test was performed to calculate significance (p<0.01 = significant; n.s. = nonsignificant). B) CCNB1 degradation rates around anaphase onset in indicated cell lines (from experiment shown in A.). The slope of a linear fit for the measured decrease in CCNB1-mCerulean3 intensities around anaphase onset is plotted for individual cells. The number of analysed cells is given as n. The table shows the mean CCNB1 degradation rate and its standard deviation. The median CCNB1 degradation rate is drawn as a black line. A two-tailed t-test was performed to calculate significance (p<0.01 = significant; n.s. = non-significant)). C) SILAC ratios for APC/C and MCC subunits detected in mass spectrometry analysis of APC8-mCherry pulldown with or without SAC activation. APC/C was purified via APC8-mCherry pulldowns from untreated cells (medium SILAC condition) and cells treated with 20 nM nocodazole (noc) for 18 hours (heavy SILAC condition). Depicted are the combined SILAC ratios from three technical replicates. The light SILAC condition was used as a reference for unspecific binding to the affinity beads. D) SILAC ratios for APC/C and MCC subunits detected in mass spectrometry analysis of APC8-mCherry pulldown from APC8-mCherry ΔAPC7 cells as described in C). E) Indicated cell lines were treated with control siRNA or MAD2 targeting siRNA for 96 hours, stained with Hoechst and imaged. Representative images from one of the three independent experiments are shown. F) Quantification of normal nuclei from experiments described in E). From each experiment at least 500 cells were analysed per condition and the cumulative percentage of normal nuclei from all three experiments is shown.

Overall, these results indicate that loss of APC7 or APC16 has little impact on mitotic progression, APC/C activity towards CCNB1, and SAC functionality. However, cell lines lacking either APC7 or APC16 appear to rely less on MAD2 for maintaining their genome stability.

### Genetic ablation of Mad2 in cells deleted of APC7 or APC16

As ΔAPC7 and ΔAPC16 cell lines displayed reduced sensitivity to MAD2 depletion (Figure 2F), we sought to test whether genetic ablation of MAD2 is tolerated in these cell lines. To assess the synthetic viability of MAD2 deletion with the loss of APC7, we mixed equal numbers (i.e. 1:1:1) of wild-type, APC8-mCherry, and APC8-mCherry ΔAPC7 cells (Figure 3A, Figure S3A) and performed CRISPR-mediated MAD2 deletion in the resulting mixed cell population. After selection, single colonies were analysed for the absence of MAD2 by immunoblotting. We recovered a total of six AMAD2 cell lines, all of which were derived from the APC8-mCherry ΔAPC7 background (Figure 3B). This result is consistent with MAD2 being essential in human cells and demonstrates that cells lacking APC7 can tolerate loss of MAD2. To confirm the observed synthetic viability and assess whether it also applies to APC16 deletion, we devised an approach facilitating direct deletion of MAD2 from wild type cells along with co-deletion of either APC7 or APC16. For this we GFP-tagged MAD2 endogenously and transfected these cells with a MAD2 deletion construct bearing a selection marker together with a selection-less plasmid encoding a guide RNA targeting either APC7 or APC16 (Figure 3C, Figure S3B). After selection, clonal cell colonies were isolated based on the visually observed loss of GFP fluorescence (indicating loss of MAD2-GFP) and subsequently analysed by immunoblotting for the absence of MAD2-GFP. Concurrent loss of APC7 and APC16 was assessed by immunoblotting and genotyping, respectively. Using this approach, we obtained four AMAD2-GFPΔAPC7 and six AMAD2-GFPΔAPC16 cell lines (Figure 3D, Figure S3C, Figure S3D). As the experimental setup only selects for MAD2 deletion, the strict co-occurrence of either APC7 or APC16 loss with MAD2 deletion indicates that the observed viability is synthetic. Consistent with the deletion of MAD2 and concurrent loss of SAC activity, neither ΔAPC7AMAD2 nor AMAD2-GFPΔAPC16 cells arrested in mitosis upon nocodazole treatment (Figure 3E). Taken together, these results show that genetic deletion of either APC7 or APC16 allows for continued proliferation in absence of MAD2.

**Figure 3.**
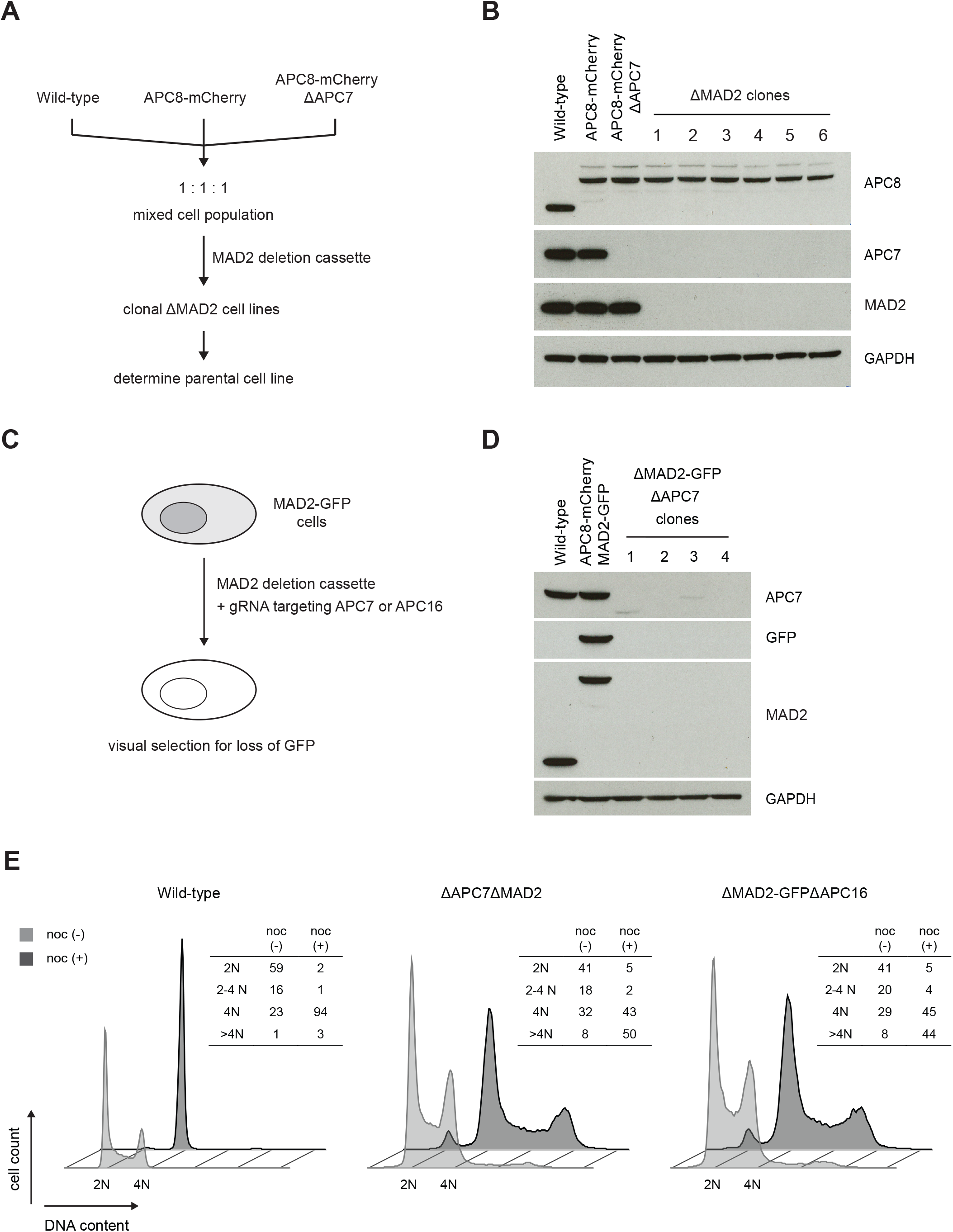
Loss of APC7 or APC16 provides synthetic viability to MAD2 deletion. A) Strategy for assessing the synthetic viability of MAD2 deletion with the ΔAPC7 genotype. An equal number of wild-type, APC8-mCherry and APC8-mCherry ΔAPC7 cells were mixed and seeded together, resulting in a mixed cell population with one-third of cells lacking APC7. The genomic MAD2 locus was then targeted by Crispr/Cas9 using a selectable donor plasmid designed to disrupt the MAD2 gene. After selection, each clonal cell line was analysed by immunoblotting to assess loss of MAD2 and to determine its parental cell line. B) Analysis of APC7 and MAD2 expression in the six clonal AMAD2 cell lines retrieved from the setup outlined in A). GAPDH levels were analysed as a loading control. Note that all six retrieved clonal cell lines derive from APC8-mCherry ΔAPC7 cells. C) Scheme of Crispr/Cas9-based MAD2-GFP synthetic viability assay. Cells expressing endogenously GFP-tagged MAD2 (MAD2-GFP) were transfected with a selectable deletion cassette for the MAD2 gene along with a plasmid encoding a guide RNA targeting either APC7 or APC16. After selection, cell colonies were microscopically inspected for loss of MAD2-GFP fluorescence. Cell colonies lacking green fluorescence were then analysed by immunoblotting for loss of MAD2 expression. The status of APC7 and APC16 was assessed by immunoblotting or sequencing of the genomic locus, respectively. D) Immunoblot analysis of the four clones retrieved from the MAD2-GFP synthetic viability assay performed in combination with an APC7 targeting guide RNA. Note that all retrieved clonal cell lines lost expression of APC7. E) Analysis of SAC functionality in ΔAPC7ΔMAD2 and ΔMAD2-GFPΔAPC16 cells. Cellular DNA from wild-type, ΔAPC7ΔMAD2 and ΔMAD2-GFPΔAPC16 cells, with or without 18 hours of 20 nM nocodazole, was stained with propidium iodide and analysed by FACS. The tables show the percentage of cells with the respective (2N, 2N-4N, 4N and >4N) DNA content.

### Analysis of cells lacking MAD2

Loss of SAC activity leads to genomic instability (Dobles et al. 2000, Kops et al. 2005 and Figure 2E, F). We therefore analysed the cellular DNA content of different ΔAPC7ΔMAD2 and ΔMAD2-GFPΔAPC16 clonal cell lines by FACS (Figure S4A). A modest increase in polyploid cells was observed in the SAC deficient cell lines compared to wild-type cells, indicating ongoing, yet tolerable chromosomal instability. To further explore the genomic stability of ΔAPC7ΔMAD2 cells, we compared mitosis and chromosome segregation in ΔAPC7ΔMAD2 cells with wild-type cells and wild-type cells depleted of MAD2 by RNAi. To this end, we fluorescently tagged histone H2B in wild-type and ΔAPC7ΔMAD2 cells and analyzed these cells by live cell imaging (Figure 4A). We observed a decreased time from chromatin condensation to anaphase onset in ΔAPC7ΔMAD2 cells compared to wild type cells, consistent with MAD2 causing delayed mitotic progression in wild-type cells (Figure 4B). Wild-type cells depleted of MAD2 by RNAi most frequently (85%) proceed through mitosis without formation of a metaphase plate, resulting in aberrant or undetectable chromosome segregation (Figure 4A, Figure 4C and Figure S4B). In contrast, ΔAPC7ΔMAD2 cells frequently (96%) formed a metaphase plate despite lacking MAD2 function and segregated chromosomes (Figure 4A and Figure 4C). Compared to wild-type cells, we captured more chromosome segregation errors in ΔAPC7ΔMAD2 cells in our live cell imaging data (Figure 4C and Figure S4B), indicating an increased rate of ongoing segregation errors. As MPS1 acts upstream of MAD2 in the SAC response, we tested whether genetic ablation of MAD2 would relieve cells from the need for MPS1 kinase activity to maintain their genome stability. FACS analysis of ΔAPC7ΔMAD2 or ΔMAD2-GFPΔAPC16 cells upon treatment with the MPS1 inhibitor reversine shows a marked increase in cells will >4N DNA content (Figure 4D). Hence, MAD2 deletion does not relieve these cell lines from the need for MPS1 activity to prevent excessive polyploidy. In summary, cells lacking MAD2 and APC7 display an accelerated mitosis concurrent with increased chromosome segregation errors and require MPS1 activity for genome stability in absence of a functional SAC.

**Figure 4.**
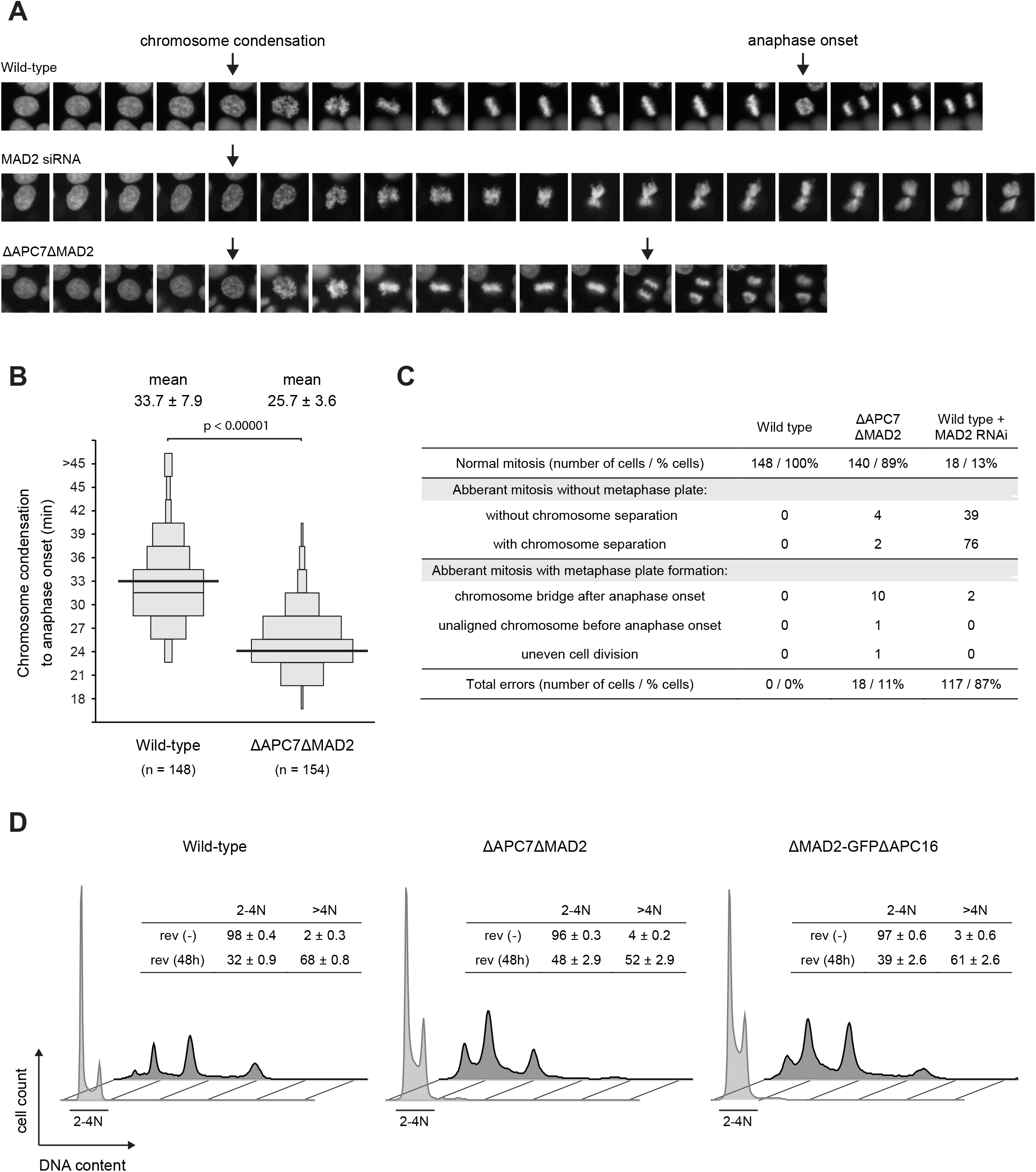
Analysis of mitosis in ΔAPC7AMAD2 cells. A) Representative images of mitosis observed for indicated cell lines expressing H2B-mCerulean3. The frames scored as chromosome condensation and anaphase onset are indicated with the first and second arrow, respectively. Note that no anaphase onset was scored for cells that failed to form a metaphase plate in the MAD2 RNAi condition. B) Quantification of timing from chromatin condensation to anaphase onset from experiment shown in A). Cells with identical timing are illustrated as a scaled box (as described in Figure 2A). Mean timing and the standard deviation are stated on top, median timing is drawn as a black line. Results of two independent experiments are shown and the number of analysed cells is stated as n. A two-tailed t-test was performed to calculate significance. C) Table summarizing observed chromosome segregation errors in indicated cell lines and conditions. D) FACS-profiles of ΔMAD2ΔAPC7 and ΔMAD2-GFPΔAPC16 cells treated with reversine. Cellular DNA from wild-type, ΔAPC7ΔMAD2 and ΔMAD2-GFPΔAPC16 cells, with or without 48 hours of 0.5 |aM reversine, was stained with propidium iodide and analysed by FACS. The table shows the percentage of cells with the respective (2N-4N and >4N) DNA content.

## Discussion

The APC/C complex, composed of 19 subunits, is the largest known ubiquitin ligase. How this large complex assembles in vivo is not fully understood. Here we have deleted the two non-essential, higher eukaryote-specific APC/C subunits APC7 and APC16, and analysed their role in APC/C assembly. This analysis revealed that APC16 is required for APC7 assembly into the APC/C in vivo, and that APC16 can incorporate into the APC/C independently of APC7. Gratifyingly, these results are in line with in vitro data on an APC3/APC7/APC16 subcomplex (Yamaguchi et al. 2015). Prior to high resolution information about APC/C structure, a systematic deletion of APC/C subunits in yeast identified APC/C sub-complexes, which may constitute potential building blocks during APC/C assembly (Thornton et al. 2006). Similarly, the analysis of APC/C assembly upon deletion of individual APC/C subunits may offer an approach to delineate an in vivo APC/C assembly path in human cells and uncover assembly or chaperoning factors involved. Such a strategy will, however, require conditional APC/C subunit deletions (or depletions), as the majority of APC/C subunits is essential for cell viability.

Analysis of key aspects of APC/C function, namely mitotic timing, CCNB1 degradation and response to spindle assembly defects, revealed no significant alterations upon loss of either APC7 or APC16. However, depletion of APC16 by RNAi in human cells and *C. elegans* results in mitotic phenotypes consistent with impaired APC/C function (Kops et al. 2010). It is currently unclear why APC16 depletion and APC16 deletion result in different phenotypes, but the difference between acute and permanent loss may contribute to this discrepancy. The sole, yet striking, phenotype detected in our study for cells lacking either APC7 or APC16 is their reduced dependence on MAD2 function for viability, indirectly revealing a functional impact of these APC/C subunits in HCT116 cells. The penetrance of genome instability upon MAD2 depletion by RNAi is significantly reduced in ΔAPC7 and ΔAPC16 cells compared to wild-type HCT116 cells. This reduced dependence on MAD2 function ultimately allows for genetic ablation of MAD2 in ΔAPC7 and ΔAPC16 cell lines. The synthetic viability between loss of MAD2 and deletion of either APC7 or APC16 may be particularly notable given the lack of a pronounced increase in mitotic length in ΔAPC7 and ΔAPC16 cells. Our previous work showed that deletion of two APC/C-employed E2 enzymes, UBE2S and UBE2C, provides viability to loss of MAD2 (Wild et al. 2016). ΔUBE2SΔUBE2C cells display a significantly prolonged mitosis, and it was therefore rationalized that it is this increased length of mitosis which enables cells to properly segregate the genomic material in absence of SAC activity. In contrast, mitosis was not detectably prolonged in ΔAPC7 and ΔAPC16 cells, yet these genetic backgrounds provide synthetic viability to MAD2 deletion. Therefore, even a mild mitotic delay, which may be below the detection limits of our live cell assay, may suffice to render MAD2 function non-essential in ΔAPC7 and ΔAPC16 cells. Alternatively, other, yet to be discovered, mechanisms to sustain genome stability in the absence of SAC signalling may exist. In the context of this study we focused on the analysis of mitotic APC/C activity, yet other aspects of APC/C function have been reported and their dependence on either APC7 or APC16 remains to be determined (Harper et al. 2002, Eguren et al. 2011).

Deletion of MAD2 has been shown to be embryonically lethal in mice, and MAD2 haplo-insufficiency significantly impairs SAC function and accurate chromosome segregation in both murine and human cells (Dobles et al. 2000, Michel et al. 2001). The first MAD2 deletion from murine cells was reported in the context of concurrent TP53 deletion, possibly reflecting an increased tolerance of TP53 null cells towards aneuploidy (Burds et al. 2005). Subsequently, it was discovered that mice lacking MAD2 in specific cell types survive, albeit with an increased rate of aneuploidy (Foijer et al. 2013) or onset of ultimately lethal diseases (Foijer et al. 2017). It will be of interest to dissect the distinct properties of reported MAD2 knockout cells in terms of their chromosome segregation error rates on the one hand and potentially altered tolerance towards aneuploidy on the other hand.

The kinase MPS1 has established functions outside of SAC signaling (Liu and Winey, 2012). We find that cells lacking MAD2, and thus SAC function, still require MPS1 for maintaining their genome stability. Hence, SAC-independent MPS1 functions, such as its role in chromosome alignment (Jelluma et al. 2008), are essential for genome integrity. This scenario is reminiscent of yeast, an organism in which Mad2 is not essential for viability, yet Mps1 is. Our genetic MAD2 knockout cells could provide a useful tool to further elucidate SAC-independent, essential functions of MPS1.

Both MPS1 and MAD2 act to maintain the stable euploid state of a cellular genome. The contribution of genomic instability to human disease has been of great research interest ever since the initial observation that aneuploidy is commonly detected in human cancers (Boveri 1902, Kops et al. 2005, Funk et al. 2016). We envision that further genetic interrogations of cellular mechanisms controlling genome stability, such as the SAC, will significantly advance our understanding of mechanisms preventing genome instability on the one side, and mechanisms underlying an increased tolerance towards ongoing genome instability on the other side.

## Methods and Materials

### FACS

To analyse the distribution of cellular DNA content, exponentially growing cells were harvested at ~70 % confluency using trypsin. Cells were pelleted, resuspended in 500 μl ice-cold PBS and then permeablized by addition of 500 μl ice-cold ethanol. Samples were incubated on ice for 45 min, pelleted by centrifugation and washed with cold PBS. The cell pellet was resuspended in propidium iodide (PI) staining solution (10 μg/ml PI in PBS) supplemented with RNAse (25 μg/ml) and incubated for 30 min at 37 °C. Samples were analyzed by flow cytometry using a BD LSR Fortessa (BD Biosciences). Data were analyzed and visualized using the FlowJo software (FlowJo, LLC). The Cell Cycle function in FlowJo was used to quantify cell populations with different DNA content or, in case of perturbation experiments, gates were manually set for one sample and the defined gates then applied to all samples of the same experiment.

### APC/C affinity purification

APC/C was purified from APC8-mCherry expressing cells using RFP-Trap (RFP-Trap^®^_MA, ChromoTek). For SILAC-based quantification of proteins (Ong et al. 2002), cells were grown in SILAC medium supplemented with natural variants of L-arginine (Arg) and L-lysine (Lys), i.e. Arg^0^/Lys^0^ for the light SILAC condition, or supplemented with isotope labeled variants of Arg and Lys, i.e. Arg^6^/Lys^4^ for the medium and Arg^10^/Lys^8^ for the heavy SILAC condition. For APC/C purification, cells were lysed in RIPA buffer (50 mM Tris-HCl pH 7.5, 150 mM NaCl, 1% Nonidet P-40, 0.1% sodium-dodecyl-sulfate, 1 mM EDTA) supplemented with protease inhibitors (Complete protease inhibitor mixture tablets, Roche Diagnostics). Lysates were incubated for 10 min on ice and cleared by centrifugation at 16,000 × g. Cleared lysates were then incubated on a rotating wheel for 1 hour at 4 °C with 20 μl of RFP-Trap beads. Beads were washed 3 times with RIPA buffer and eluted with LDS sample buffer (Thermo Fisher Scientific). For SILAC experiments, equal amount of proteins, as determined by Bradford assay (Bio-Rad), were incubated with the RFP-Trap beads. The beads from the different SILAC conditions were washed once separately with RIPA buffer, combined into one new tube and subsequently washed together 3 times with RIPA buffer. Eluates were analysed by SDS-PAGE followed by immunoblotting or mass spectrometry.

### Mass spectrometry

Samples in LDS sample buffer were incubated with dithiothreitol (10 mM) for 10 min at 70 °C and subsequently with chloroacetamide (5.5 mM) for 60 min at 25 °C. Proteins were separated on a 4%–12% gradient SDS-PAGE, stained with colloidal coomassie blue, and digested using in-gel digestion method (Jensen et al. 1999). In brief, gel lanes were cut into six fractions and each gel piece was further sliced into smaller pieces (~1mm). Gel pieces were destained with 50% ethanol in 25 mM ammonium bicarbonate (pH 8.0) on a rotating wheel at room temperature and dehydrated with 100% ethanol. Trypsin (in 25 mM ammonium bicarbonate pH 8.0) was added to the dry gel pieces and incubated overnight at 37 °C. The trypsin digestion was stopped by addition of trifluoroacetic acid (0.5% final concentration) and peptides were extracted from the gel pieces by stepwise increase in acetonitrile concentration (to 100% final). Next, acetonitrile was removed by centrifugal evaporation and peptides were purified by a C18 reversed-phase packed Stage-Tip. Peptide fractions eluted from Stage-Tips were analyzed on a quadrupole Orbitrap (Q-Exactive, Thermo Scientific) mass spectrometer equipped with a nanoflow HPLC system (Thermo Scientific). Peptide samples were loaded onto C18 reversed-phase columns and eluted with a linear gradient from 8 to 40% acetonitrile containing 0.5% acetic acid in 105 min gradient. The Q-Exactive was operated in the data dependent mode automatically switching between single-mass-spectrometry and tandem-mass-spectrometry acquisition. Survey full-scan MS spectra (m/z 300–1700) were acquired in the Orbitrap. The 10 most intense ions were sequentially isolated and fragmented by higher-energy C-trap dissociation (HCD). Peptides with unassigned charge states, as well as peptides with charge state less than +2 were excluded from fragmentation. Fragment spectra were acquired in the Orbitrap mass analyzer.

Raw MS data were analyzed by the MaxQuant software (Cox and Mann 2008). Mass spectra were searched against protein sequences from the UniProt knowledge base using the Andromeda search engine (Cox et al. 2011). Spectra were searched with a mass tolerance of 6 ppm for precursor ions, 20 ppm for fragment ions, strict trypsin specificity and allowing up to two missed cleavage sites. Cysteine carbamido methylation was searched as a fixed modification, whereas amino-terminal protein acetylation and methionine oxidation were searched as variable modifications. A false discovery rate of less than one percent was achieved using target-decoy search strategy (Elias and Gygi 2007) and a posterior error probability filter.

### Antibodies

α-MAD2 (EMD Millipore, MABE866), α-APC1 (Cell Signaling Technology, APC1 (D1E9D) Rabbit mAb #13329), α-APC2 (Cell Signaling Technology, APC2 Antibody #12301), α-APC7 (Bethyl laboratories, A302-551A), α-APC8 (Cell Signaling Technology, APC8 (D5O2D) Rabbit mAb #15100), α-APC11 (Cell Signaling Technology, APC11 (D1E7Q) Rabbit mAb #14090), α-GAPDH (Merck Millipore, Anti-GAPDH Antibody, ABS16), α-GFP (Cell Signaling Technology, GFP (D5.1) XP^®^ Rabbit mAb), α-CCNB1 (BD Pharmingen, Purified Mouse Anti-Cyclin B1, Clone GNS-1)

### Genome engineering

Crispr/Cas9 was applied to tag or delete genes of interest. DNA oligonucleotides encoding the guide RNAs were cloned into pX330 (Cong et al.) and co-transfected with donor plasmids to achieve the desired modification within the genome of HCT116 cells. Donor plasmids generally contain homology arms spanning approximately 500 bp from each side of the cutting site (as designated by the guide RNAs) and a selection marker, providing resistance towards puromycin, G418, zeocin, hygromycin, blasticidin or NTC (Kochupurakkal and Iglehart). For tagging, mCherry, EGFP, mVenus or mCerulean3 (Markwardt et al.) were fused in frame to the last exon of the gene of interest. After selection, clonal cell lines were analysed by genomic PCR and/or immunoblotting to screen for the desired genomic modification. The following guide RNAs (including PAM motif shown in bold) were used (when two guide RNAs are listed both were used simultaneously):

For gene tagging:

APC8: **CCA**TAGTTGGCTACTCTCAAGCC

MAD2: aacacaatcactaaattgcacgg and tgaccttttccagcagtgagtgg

CCNB1: TGTAACTTGTAAACTTGAGTTGG

H2B: **CCA**CGCAtgttttcaataaatga and aatcatttcattcaaaagggggg

For gene deletions:

dAPC7: TGCGGGACATGGCGGCCGCGGGG

dAPC7intron: agcgcgactgtcacatcgctagg and agctcagggacccagcctcctgg

dAPC16: GCCCTTTTCACCTACCCCAAAGG

dMAD2: **CCC**TGCGCGGGAGCGCCGAAATC

The following table summarizes the cell lines generated in this study:

| no. | cell line | parental line | additional resistance* |
| --- | --- | --- | --- |
| 1 | Apc8-mCherry | HCT116 | G418 |
| 2 | Apc8-mCherry ΔAPC7 | no. 1 | hygromycin / zeocin |
| 3 | Apc8-mCherry ΔAPC16 | no. 1 | hygromycin / zeocin |
| 4 | CCNB1-mCerulean3 | HCT116 | blasticidin |
| 5 | CCNB1-mCerulean3 H2BD-mVenus | no. 4 | NTC |
| 6 | CCNB1-mCerulean3 ΔAPC7 | no. 4 | hygromycin / zeocin |
| 7 | CCNB1-mCerulean3 ΔAPC7 H2BD-mVenus | no. 6 | NTC |
| 8 | CCNB1-mCerulean3 ΔAPC16 | no. 4 | hygromycin / zeocin |
| 9 | CCNB1-mCerulean3 ΔAPC16 H2BD-mVenus | no. 8 | NTC |
| 10 | Apc8-mCherry ΔAPC7 ΔMad2 | no. 2 | puromycin / blasticidin |
| 11 | Apc8-mCherry Mad2-EGFP | no. 1 | puromycin |
| 12 | Apc8-mCherry ΔMad2-EGFP ΔAPC7 | no. 11 | hygromycin |
| 13 | Apc8-mCherry ΔMad2-EGFP ΔAPC16 | no. 11 | hygromycin |
| 14 | H2BD-mCerulean3 | HCT116 | blasticidin |
| 15 | Apc8-mCherry ΔAPC7 ΔMad2 H2BD-mCerulean3 | no. 10 | NTC |

\*if stated, the two selection markers were used simultaneously (offering a stringent selection regime for targeting both alleles of the gene of interest)

### Live cell microscopy

Cells were grown in 24 well plates suited for microscopy (Imaging Plate CG, zell-kontakt). For live cell imaging, cells were seeded into FluoroBrite DMEM medium (Thermo Fisher Scientific) supplemented with 10% FBS and 4 mM L-glutamine. Live cell imaging was performed on an Olympus ScanR microscope using a 20 × objective. Images were taken continuously every 3 min acquiring 6 z-stacks of 1.5 μm (H2B-mCerulean3 expressing cell lines) or every 5 min acquiring 4 z-stacks of 2 μm (CCNB1-mCerulean3 and H2B-mVenus expressing cell lines).

### RNAi

Cells were seeded into a 24-well plate and transfected with siRNAs 24 hours later using RNAiMAX (Lipofectamine^®^) as transfection reagent according to the manufactures guidelines. For MAD2 depletion, the following siRNA was used: CGCCUUCGUUCAUUUACUAtt (Silencer Select, Thermo Fisher Scientific). Silencer Select Negative Control #1 siRNA (s813, Thermo Fisher Scientific) was used as a negative control. The final concentration of siRNAs was 5 nM. 24 hours after siRNA transfection, cells were trypsinized and reseeded into 24-well plates suitable for microscopy (Imaging Plate CG, zell-kontakt). 72 hours later (96 hours after siRNA transfection), Hoechst (Hoechst 33342, Life Technologies, H3570) was added directly to the medium and cells imaged with an Olympus ScanR microscope using a 20 x objective. After imaging, cell lysates were prepared from the imaged 24-wells using RIPA buffer (50 mM Tris-HCl pH 7.5, 150 mM NaCl, 1% Nonidet P-40, 0.1% sodium-dodecyl-sulfate, 1 mM EDTA). Samples containing equal amounts of protein, as determined by Bradford assay (Bio-Rad), were mixed with LDS-sample buffer and analysed by SDS-PAGE followed by immunoblotting. For live cell imaging, cells were transfected with MAD2 siRNA/RNAiMAX mix 32 hours prior to imaging.

### Image analysis

Images acquired by live cell microscopy were analysed using the Imaris Image analysis software (Bitplane). Nuclear CCNB1 influx was visually scored as the first frame prior to mitosis in which nuclear CCNB1-mCerulean3 intensity equalled or exceeded cytoplasmic CCNB1-mCerulean3 intensity. Anaphase onset was scored as the first frame in which two separate chromosome masses were observed in the H2B-mVenus channel after metaphase plate formation. To obtain CCNB1 degradation rates, z-projections (median projections) were generated in Fiji and chromatin objects (Surfaces) were generated in Imaris using the Magic Wand tool on the H2B-mVenus channel. Mean CCNB1-Cerulean3 intensities of the defined chromatin objects were extracted to obtain CCNB1 degradation trajectories for individual cells. The CCNB1 degradation rate at anaphase onset was calculated as the slope of the linear fit for the mean CCNB1 intensities one frame before anaphase onset, at anaphase onset and one frame after anaphase onset. For H2B-mCerulean3 expressing cells, mitotic timing was calculated from the onset of chromosome condensation to anaphase onset. To score chromosome segregation errors, each analysed cell was visually scanned through the six acquired z-stacks for observable chromosome segregation defects around anaphase onset.

To analyse nuclear size and shape upon Mad2 depletion, images of Hoechst stained nuclei were visually separated into two categories: normally sized and shaped nuclei (i.e. normal nuclei), and enlarged and irregularly shaped nuclei (i.e. aberrant nuclei).

### Genomic analysis of APC16 locus

The genomic locus of APC16 encompassing the sequence targeted by the guide RNA was analysed with the following primer set: gAPC16fw (5′-gcgcGGTACCgttcaaaacctgactgatattttggcctgagac) and gAPC16rev (5′-gcgcCTTAAGcgtgcccagccctgactctgcctttaacctgg). KOD Xtreme Hot Start DNA Polymerase (Novagen, Toyobo) was used according to the manufactures protocol to amplify the APC16 locus. GeneRuler 1 kb DNA Ladder (Thermo Fisher Scientific) was used as a DNA marker. To sequence potential indels within the APC16 locus, the PCR product was cloned into pcDNA3.1/ZEO (+) (Thermo Fisher Scientific, Invitrogen) using the KpnI and AflII restriction sites.

### Drug treatments

To assess spindle assembly checkpoint functionality, cells were treated with 200 nM nocodazole (Sigma Aldrich, M1404) for indicated times. To assay for the cellular dependence on MPS1 kinase activity, cells were treated with 0.5 μm reversine (MedChemexpress, HY-14711) for indicated times.

### Plasmids and transfections

The coding sequence of APC16 was cloned NheI/BamHI into pEGFP-N2 (Clontech), and the resulting construct (pAPC16-EGFP) was verified by sequencing. For plasmid transfection, 5 μg of plasmid (pAPC16-EGFP) was incubated in DMEM medium (without any supplements) with 15 ul TurboFect (Thermo Fisher) for 20 min at room temperature and then added to 50% confluent cells in a 10 cm dish. APC/C affinity purification was performed 32 hours after transfection.

## Acknowledgements

We thank the members of our group for their help and discussions. We thank Rajat Gupta for sharing plasmids and Elina Maskey for providing excellent technical assistance. CC is supported by the Hallas Møller Investigator Fellowship from the Novo Nordisk Foundation (NNF14OC0008541). This project has received funding from the European Research Council (ERC) under the European Union’s Horizon 2020 research and innovation program (grant agreement No 648039). We thank Dr. Simone Schopper from the CPR Mass Spectrometry Platform for her assistance with maintenance and support of the platform. We thank Gelo dela Cruz from the Flow Cytometry Core Facility at the NNF Center for Stem Cell Biology (DanStem) and Jutta Bulkescher from the CPR Protein Imaging Platform for their technical assistance with flow cytometry and microscopy, respectively. The Center for Chromosome Stability is supported by the Danish National Research Foundation. The Novo Nordisk Foundation Center for Protein Research is supported financially by the Novo Nordisk Foundation (Grant agreement: NNF14CC0001).

## Author contributions

TW and MB designed and performed the experiments, and analyzed the data. GK helped with analysis of live cell imaging data. TW, MB and CC wrote the manuscript. CC supervised the research. All authors read and commented on the manuscript.

---

**Supplemental figure 1.**
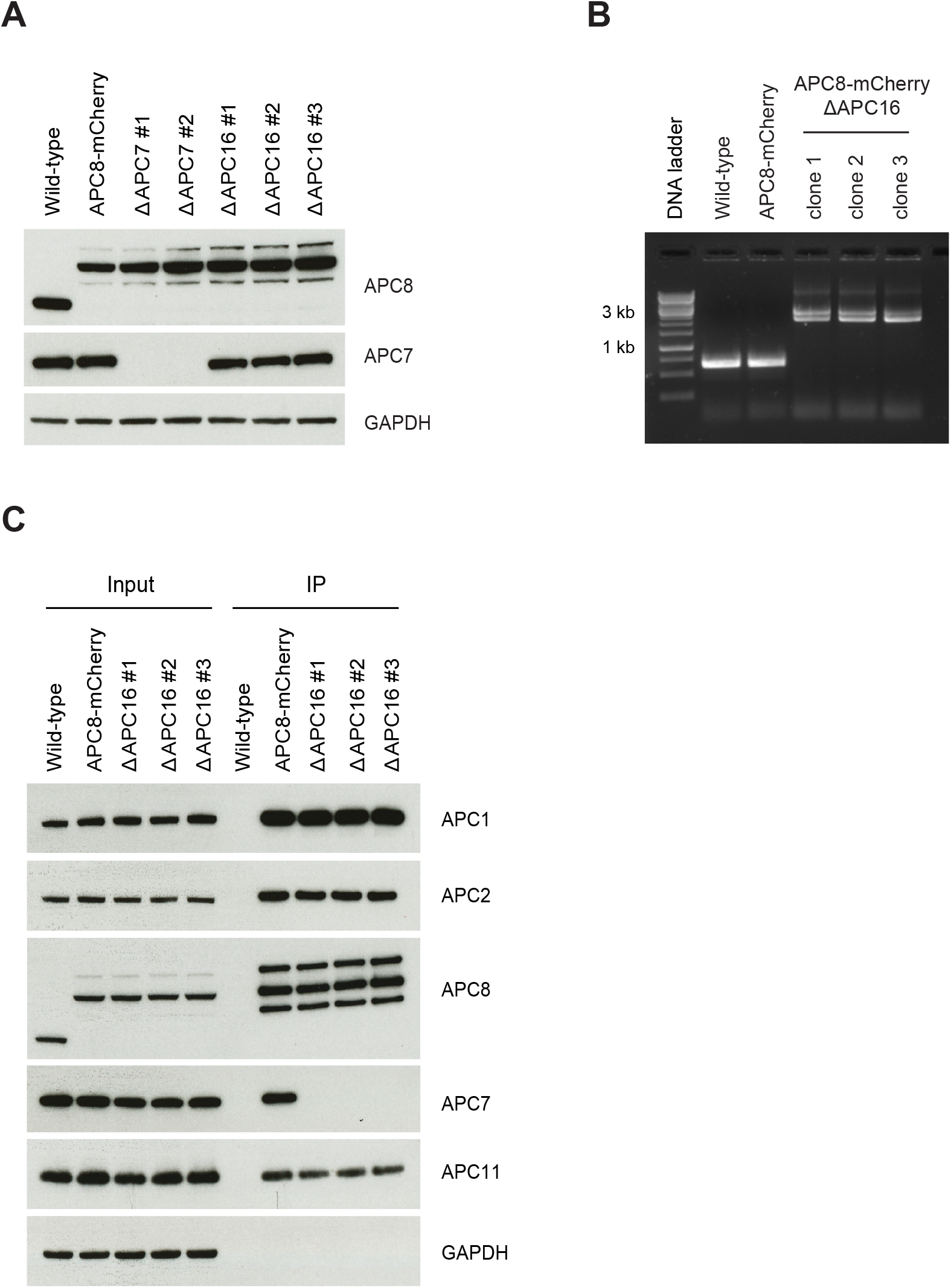
A) Cell lines generated to analyse APC/C composition upon deletion of either APC7 or APC16. Immunoblot analysis of wild-type, APC8-mCherry, APC8-mCherry ΔAPC7 and APC8-mCherry ΔAPC16 cell lines using indicated antibodies. Two independent ΔAPC7 and three independent ΔAPC16 clonal cell lines are shown. GAPDH levels were analysed as a loading control. B) Genomic APC16 locus from wild-type, APC8-mCherry and APC8-mCherry ΔAPC16 cells analysed by PCR. The used primer pair amplifies an approximately 600 base pair (bp) product from a wild-type APC16 locus and an approximately 2300 or 2900 bp product upon genomic insertion of the deletion cassette. Note that biallelic disruption of the APC16 locus was performed using two deletion cassettes bearing different selection markers, which results in two differently sized PCR products. Three independent ΔAPC16 clonal cell lines are shown. C) Analysis of APC/C composition in ΔAPC16 cells. APC/C was purified via APC8-mCherry pulldowns from APC8-mCherry and APC8-mCherry ΔAPC16 cells (from three independent ΔAPC16 clonal cell lines) and subsequently analyzed by immunoblotting using indicated antibodies. Wild-type cells served as a control for unspecific binding to the affinity beads. GAPDH levels were analysed to verify equal amount of input for the different cell lines.

**Supplemental figure 2.**
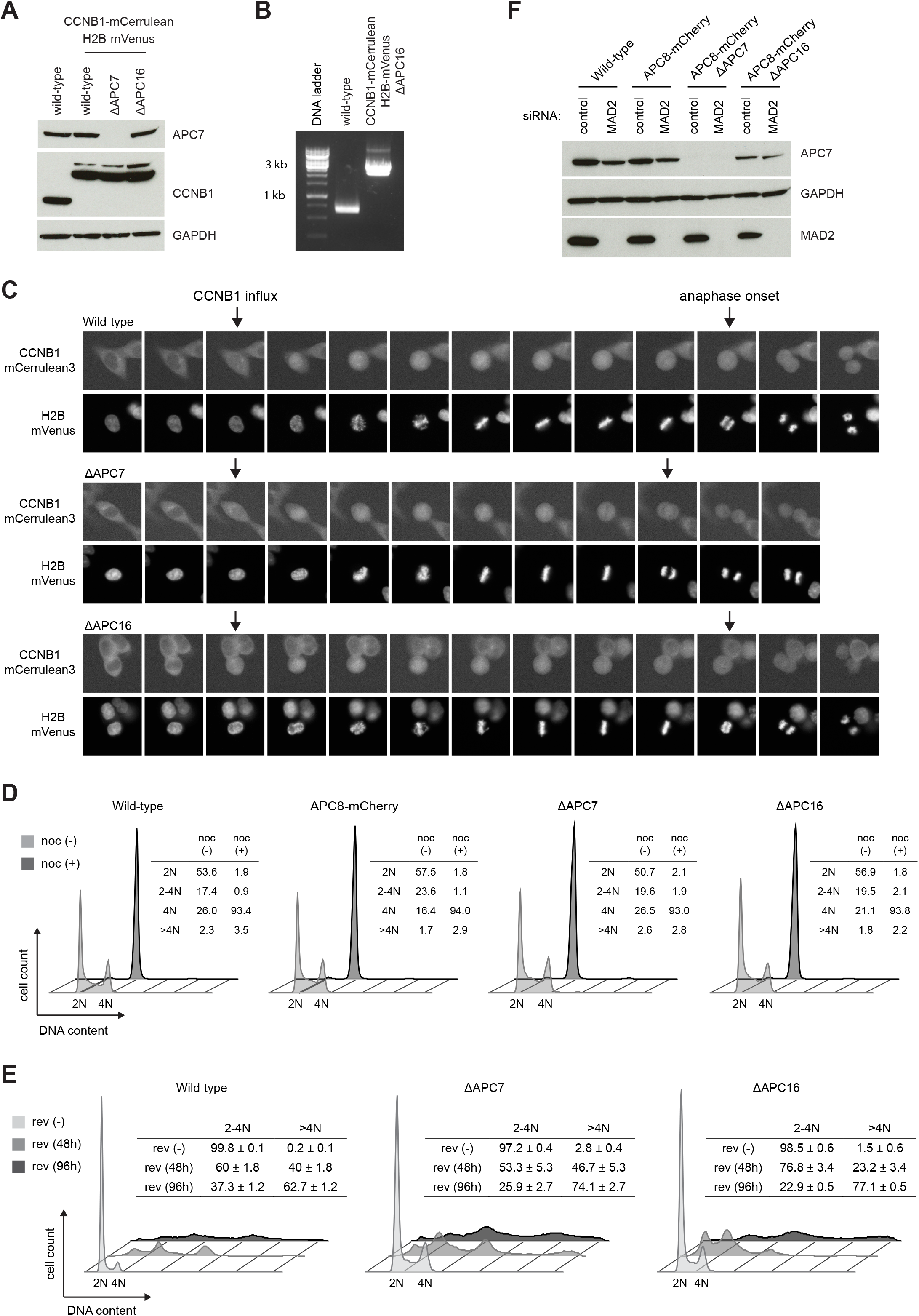
A) Immunoblot analysis of cell lines used for live cell imaging data shown in Figure 2A and 2B. Lysates from indicated cell lines were separated by SDS-PAGE, blotted and analysed with indicated antibodies. GAPDH levels were analysed to control for equal loading. B) Genomic APC16 locus from wild-type and CCNB1-mCerulean H2B-mVenus ΔAPC16 cells analysed by PCR as described in Figure S1B. C) Representative example images extracted from time-lapse imaging experiments analysed in Figure 2A and Figure 2B. For the different cell lines, the CCNB1 influx frame (CCNB1-mCerulean3 channel) and anaphase onset frame (H2B-mVenus) are indicated as the first and second arrow, respectively. D) Analysis of cellular DNA content in wild-type, ΔAPC7 and ΔAPC16 cells upon 18 hours of nocodazole treatment. The DNA content in wild-type, ΔAPC7 and ΔAPC16 cells, grown with or without 18 hours of 20 nM nocodazole, was analyzed with propidium iodide staining and flow cytometry. The tables show the percentage of cells with the respective (2N, 2N-4N, 4N and >4N) DNA content. E) The DNA content in wild-type, ΔAPC7 and ΔAPC16 cells, grown with or without 0.5 μM reversine for 48 or 96 hours, was analyzed with propidium iodide staining and flow cytometry. The histogram of one representative technical replicate is shown and the percentage of cells with 2N-4N (including both 2N and 4N) and >4N DNA content is shown in the table (percentages standard deviations were calculated from three technical replicates). F) Immunoblot analysis of control and MAD2 RNAi used in Figure 2E–F. Cells imaged for Figure 2E and Figure 2F were lysed and expression of the indicated proteins was analysed by immunoblotting. A representative blot of the three independently performed experiments is shown. GAPDH levels were analysed to control for equal loading.

**Supplemental figure 3.**
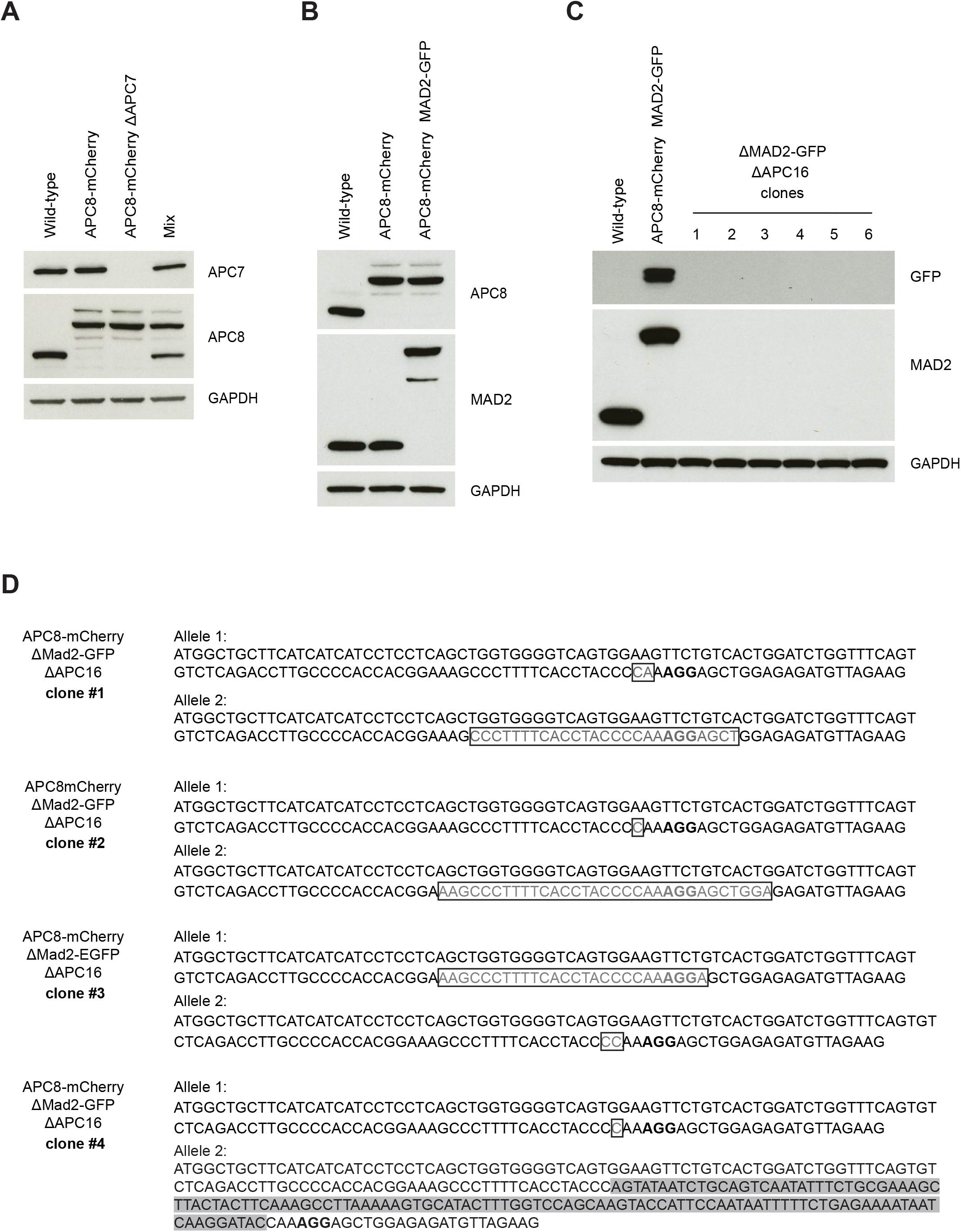
A) Immunoblot of input cells used for mixing experiment outlined in Figure 3A. Equal number of wild-type, APC8-mCherry and APC8-mCherry ΔAPC7 cells were mixed and seeded together. Lysate from the individual cell lines and the resulting mixed cell population (on the day of transfection of MAD2-targeting Crispr/Cas9 plasmids) were analysed for expression of indicated proteins with immunoblotting. B) Immunoblot analysis of the APC8-mCherry MAD2-GFP cell line along with its parental cell lines with indicated antibodies. C) Immunoblot analysis of MAD2 expression in the six clones retrieved from MAD2-GFP synthetic viability assay performed in combination with an APC16 targeting guide RNA. Immunoblot for GAPDH serves as loading control. D) Confirmation of APC16 locus disruption in four ΔMAD2-GFPΔAPC16 clones. The genomic APC16 locus surrounding the guide RNA target site was cloned from four ΔMAD2-GFPΔAPC16 clonal cell lines and sequenced. The table lists the obtained sequences, beginning with the start codon of APC16 and depicting the PAM sequence adjacent to the guide RNA in bold. Compared to the wild-type APC16 locus, deleted base pairs are shown in boxes, inserted base pairs are highlighted with grey background. Note that all sequenced mutations cause disruption of the APC16 reading frame.

**Supplemental figure 4.**
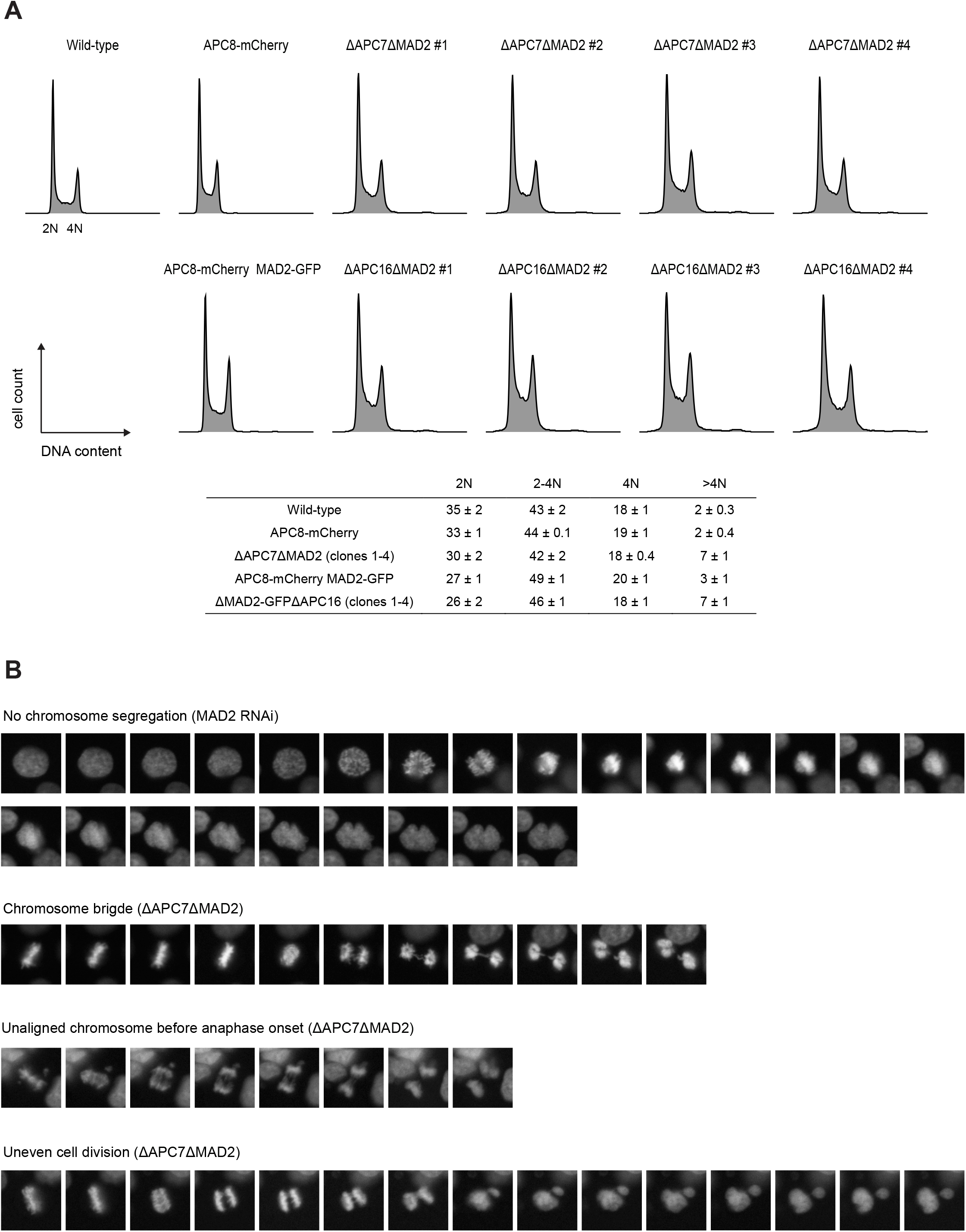
A) Analysis of DNA content in ΔAPC7ΔMAD2 and ΔMAD2-GFPΔAPC16 clonal cell lines cells. The cellular DNA from wild-type, APC8-mCherry, ΔAPC7ΔMAD2, APC8-mCherry MAD2-GFP and AMAD2-GFPΔAPC16 clonal cell lines was stained with propidium iodide and analysed by FACS. The table shows the percentage of cells with the respective (2N, 2N-4N, 4N and >4N) DNA content. B) Representative images of chromosome segregation errors summarized in Figure 4C.

